# Stable Carbon Isotopes in *Zea mays*

**DOI:** 10.1101/414789

**Authors:** Robert J. Twohey, Lucas M. Roberts, Anthony J. Studer

**Affiliations:** Department of Crop Sciences, University of Illinois Urbana-Champaign, Urbana, IL 61801, USA

**Keywords:** *Zea mays*, carbon isotopes, water-use, transpiration efficiency, vegetative phase change

## Abstract

The increasing demand for food production and predicted climate change scenarios highlight the need for improvements in crop sustainability. The efficient use of water will become increasingly important for rainfed agricultural crops even in fertile regions that have historically received ample precipitation. Improvements in water-use efficiency in *Zea mays* have been limited, and warrants a renewed effort aided by molecular breeding approaches. Progress has been constrained by the difficulty of measuring water-use in a field environment. The stable carbon isotope composition (δ^13^C) of the leaf has been proposed as an integrated signature of carbon fixation with a link to stomatal conductance. However, additional factors affecting leaf δ^13^C exist, and a limited number of studies have explored this trait in *Z. mays*. Here we present an extensive characterization of leaf δ^13^C in *Z. mays*. Significant variation in leaf δ^13^C exists across diverse lines of *Z. mays*, which we show to be heritable across several environments.

Furthermore, we examine temporal and spatial variation in leaf δ^13^C to determine the optimum sampling time to maximize the use of leaf δ^13^C as a trait. Finally, our results demonstrate the relationship between transpiration and leaf δ^13^C in the field and the greenhouse. Decreasing transpiration and soil moisture are associated with decreasing leaf δ^13^C. Taken together these results outline a strategy for using leaf δ^13^C and reveal its usefulness as a measure of transpiration efficiency under well-watered conditions rather than a predictor of performance under drought.

**Significance Statement:** This study identifies sources of variation in stable carbon isotopes of maize leaves and establishes the framework for connecting leaf δ^13^C and transpiration efficiency.

## Introduction

Increasing occurrences of extreme temperature and precipitation patterns necessitate improvements in crop productivity and sustainability (Pryor *et al.*, 2013). In the near future, abnormal and sporadic precipitation in the Midwestern United States, where agriculture is primarily rainfed, will require crops that use water efficiently. The susceptibility of *Zea mays* to yield loss due to increases in temperature and vapor pressure deficit (VPD; the difference between the amount of water in the air and the holding capacity of the air when saturated) has already been documented in this region (Lobell *et al.*, 2014). An extensive amount of time is required to develop a commercial variety with improvements in complex traits that affect canopy dynamics, photosynthetic pathways, and water-use (Hall and Richards, 2013). Thus, advancements in methods for evaluating water use are required to speed up breeding efforts to meet the imminent need for more efficient and sustainable crops.

Transpiration efficiency can be defined in agronomic terms (*TE*_*a*_ = yield ÷ transpired water) or intrinsic terms as (*TE*_*i*_ = net photosynthesis ÷ transpiration) (Dhanapal *et al.*, 2015; Ellsworth and Cousins, 2016). Transpiration is the efflux of water through stomatal pores that occurs simultaneously with the influx of CO_2_ for photosynthesis (Kim *et al.*, 2010), and can be calculated by multiplying stomatal conductance (*g*_*s*_) by VPD. Importantly, an increase in *TE* must not limit CO_2_ for photosynthesis, which would result in a reduction in yield. This is problematic in a C_3_ crop because high *g*_*s*_, which equates to high transpiration, is generally required to maintain high levels of photosynthesis. In fact, high yielding C_3_ crops have been shown to be associated with high *g*_*s*_ (reviewed in Blum, 2009). However, in C_4_ crops, the carbon concentrating mechanism maintains high concentrations of CO_2_ around Rubisco in the bundle sheath cells even when *g*_*s*_ is low (von Caemmerer, 2000). Therefore, in a C_4_ crop, there is the potential to optimize *TE*.

Direct measurements of *TE* are both time and resource consuming. These constraints are compounded by the large population size needed for a breeding program (Long and Bernacchi, 2003), but there may be high-throughput alternatives to traditional gas exchange assays. One of the proposed alternative methods is the use of carbon isotope composition of a leaf (δ^13^C, %) as an integrative measure of metabolism (von Caemmerer *et al*., 2014). Fractionation during CO_2_ uptake and assimilation processes are thought to be the major factors affecting leaf δ^13^C (Farquhar *et al.*, 1989). Post-photosynthetic fractionation, which includes all of the isotopic fractionation after Rubisco carboxylation, may also affect carbon isotope composition but its contribution to leaf δ^13^C is not known (Brüggemann et al., 2011).

The relationship between *TE* and leaf δ^13^C in C_3_ plants has made it a useful trait when breeding C_3_ crops (Farquhar *et al.*, 1984; Teulat *et al*., 2002; Rebetzke *et al*., 2002; Condon *et al.*, 2004; Saranga *et al*., 2004). In C_3_ plants, δ^13^C is negatively correlated with the ratio of leaf intercellular to ambient CO_2_ concentrations (*C*_*i*_/*C*_*a*_), which is driven mainly by the movement of CO_2_ through stomata and the rate of CO_2_ drawdown by Rubisco. As stomata close, transpiration is reduced and *C*_*i*_/*C*_*a*_ decreases. Given the theoretical relationship between *C*_*i*_/*C*_*a*_ and *TE* in C_3_ plants, there is a positive correlation between *TE* and δ^13^C (Faquhar *et al*., 1989; von Caemmerer *et al*., 2014).

The relationship between leaf δ^13^C and *TE* in C_4_ plants is more complex. Differences in isotopic fractionation between the photosynthetic types is clearly observed in the leaf δ^13^C signatures. C_3_ plants range in leaf δ^13^C from −23 to −32 ‰, whereas C_4_ plants range from −11 to −15 ‰ (O’Leary, 1988). The δ^13^C of C_4_ plants is not only related to *C*_*i*_/*C*_*a*_ but also to the proportion of CO_2_ fixed by phosphoenolpyruvate carboxylase that leaks out of the bundle sheath cells (leakiness: Φ) (Farquhar, 1983). In contrast to C_3_ plants, the relationship between δ^13^C and *C*_*i*_/*C*_*a*_ is positive except when Φ > 0.37 (Cernusak *et al.* 2013). Although Φ has been shown to be < 0.3 and stable across many environmental conditions (Henderson *et al.*, 1992), the contribution of Φ variation to observed variation in leaf δ^13^C is not completely understood. Even when Φ is constant, the variation in leaf δ^13^C in response to changes in *C*_*i*_/*C*_*a*_ is much smaller in C_4_ compared to C_3_ plants (Cernusak *et al.* 2013; von Caemmerer *et al*., 2014). These factors have limited the use of leaf δ^13^C in C_4_ plants. Additional studies that investigate variations in leaf δ^13^C and identify the underlying mechanisms that control these differences will facilitate the use of leaf δ^13^C as a proxy trait for *TE*.

*Zea mays* serves as an excellent biological system for studying photosynthesis and water use of C_4_ plants because of its economic importance as a major crop (nearly 37 million hectares planted in United States in 2017, USDA-NASS), and because of the genetic resources available. In addition, there is tremendous genomic and phenotypic diversity between lines of *Z. mays*, which is greater at the DNA sequence level than between humans and chimpanzees (Tian *et al.*, 2009). These attributes have enabled *Z. mays* to be a model system for studying a variety of complex traits through the development of populations that represent the vast intraspecies diversity (Hansey *et al.*, 2011; Romay *et al.*, 2013; Wallace *et al.*, 2014; Zhang *et al.*, 2015). Recently in a greenhouse experiment Kolbe *et al.*, (2017) assayed leaf δ^13^C of the Nested Association Mapping (NAM) founder lines, which were previously chosen to represent the genetic diversity across all maize inbreds (Flint-Garcia *et al.*, 2005; McMullen *et al*., 2009) and found substantial intraspecies variation. In this study, we focus on field-grown plants to identify factors that influence leaf δ^13^C in *Z. mays*. These results not only guide subsequent studies on leaf δ^13^C, but also provides evidence for the relationship between δ^13^C and transpiration in a C_4_ species.

## Results

### Diversity of Leaf δ^13^C in Field-Grown *Zea mays*

To assess the natural diversity of leaf δ^13^C in field grown *Z. mays*, a set of 31 inbred lines were grown for two seasons at the University of Illinois (Table S1). This set of lines included the 26 NAM founder lines, three inbreds commonly used in genetic studies and for transformation (Mo17, W22, and H99), as well as two Expired Plant Variety Protection (ExPVP) lines (LH82 and PH207). In 2015, the average leaf δ^13^C across all lines was −12.14 ‰, with a range from - 11.61 to −13.02 ‰. Slightly shifted, but similar trait values were observed in 2016, with an average leaf δ^13^C of −12.67 ‰, with a range from −12.22 to −13.29 ‰. Although there are some genotypes that depart from a linear regression of the two years of data (Figure 1), a Pearson correlation showed a significant relationship (*r* = 0.6987; *p*-value < 0.0001). To further investigate the repeatability and genetic influence on leaf δ^13^C, we incorporated data from a previous study that measured leaf δ^13^C on the NAM founder lines in a greenhouse experiment (Kolbe *et al.*, 2017). From these three different environments, we calculated broad sense heritability on a line mean basis and found leaf δ^13^C to have a moderate heritability (*H*^*2*^ = 0.57).

**Figure 1:**
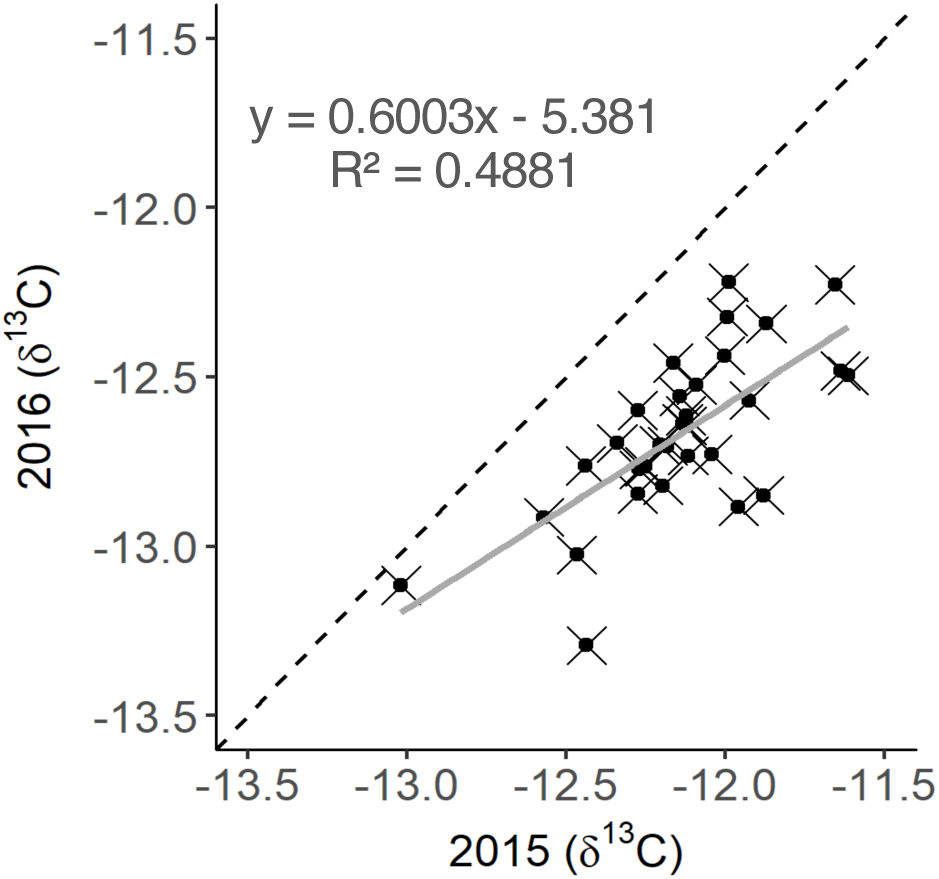
Comparison of leaf δ^13^C values from two individual field seasons. Data points represent leaf δ^13^C values measured in two consecutive years during mature growth. A linear best-fit line is shown in grey, and the dashed line shows a 1:1 relationship.

### Leaf δ^13^C Circadian Time-course

Although δ^13^C is generally considered an integrated measure of carbon fixation, the potential effect of diurnal pools of photosynthate have not previously been evaluated. To determine if leaf δ^13^C values change during a 24-hour period, we sampled the *Z. mays* reference line B73 at one-hour intervals in a circadian time-course experiment. An ANOVA was used to compare the individuals sampled across the 24-hour period and a significant difference was found between sampling times (*p* = 0.0219). A Tukey’s *post hoc* pairwise comparison revealed that this difference was due to a single time point (12:00) that was significantly lower than 5:00, 6:00, and 18:00. However, the differences do not resemble a pattern consistent with a diurnal mechanism. This result supports the use of leaf δ^13^C as an integrated measurement of carbon fixation in the leaves and not a measure of active photosynthate, which would fluctuate diurnally. The pooled samples had an average leaf δ^13^C value of −12.91 ‰ for the duration of the time-course. The maximum deviation of the pooled samples from their average was 0.24 ‰. This deviation is near expectation for technical measurement error. Samples collected and analyzed as individuals from each time point resulted in leaf δ^13^C values consistent with pooled samples (Figure 2).

**Figure 2:**
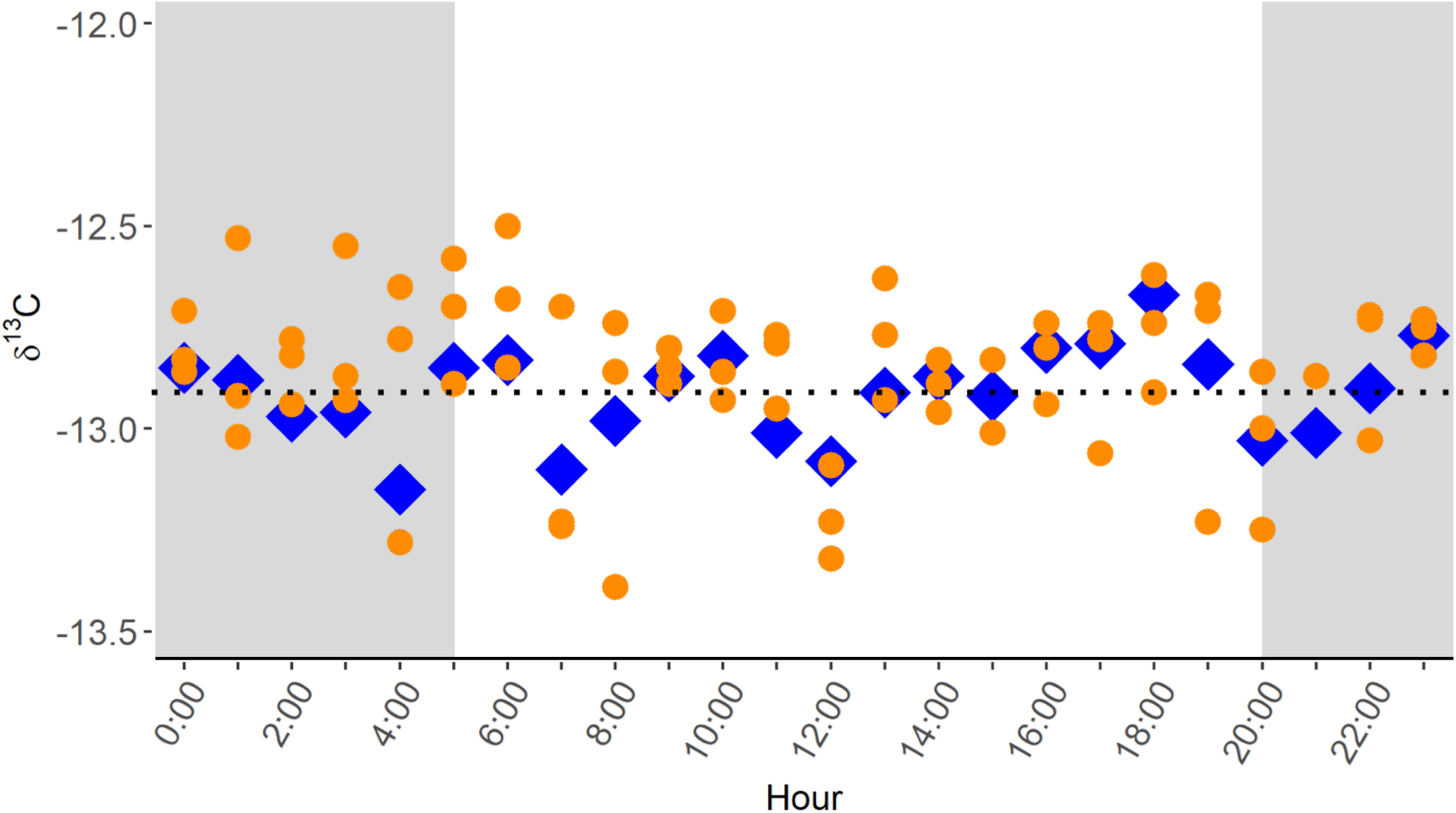
Circadian time course of leaf δ^13^C in *Z. mays*. Data points represent leaf δ^13^C values of a single *Z. mays* line (B73) collected hourly for a duration of 24 hours. Blue diamonds show leaf δ^13^C values of pooled samples and orange circles show leaf δ^13^C values of individual plants that compose each pool. The horizontal dashed line represents the average leaf δ^13^C value (−12.91 ‰) for all pooled samples during the 24 hour collection period. Shaded areas of the graph represent time of darkness between sunset and sunrise.

### Leaf δ^13^C Across *Zea mays* Development

To determine if leaf δ^13^C values change during plant growth, the *Z. mays* reference line B73 was sampled as new leaves expanded throughout development over two growing seasons. These experiments showed variation in leaf δ^13^C values across developmental time. In both years a trend emerged where leaves had less negative δ^13^C values as they transitioned between juvenile and adult leaf stages. A mixed linear model was used to combine the data from the two years so that the trend could be further examined using Least Squared Means (Figure 3A). Leaves from juvenile growth stages (V4-V5) resulted in a lower average δ^13^C value (−12.91 ‰) than adult leaves (V10-V19), which averaged −12.24 ‰. The data from both years places the transition in leaf δ^13^C values near V7 in B73. After this transition, leaf δ^13^C values stay relatively constant throughout mature growth including the flag leaf. From these results, we conclude that V10 is an optimal stage to sample leaf δ^13^C.

**Figure 3:**
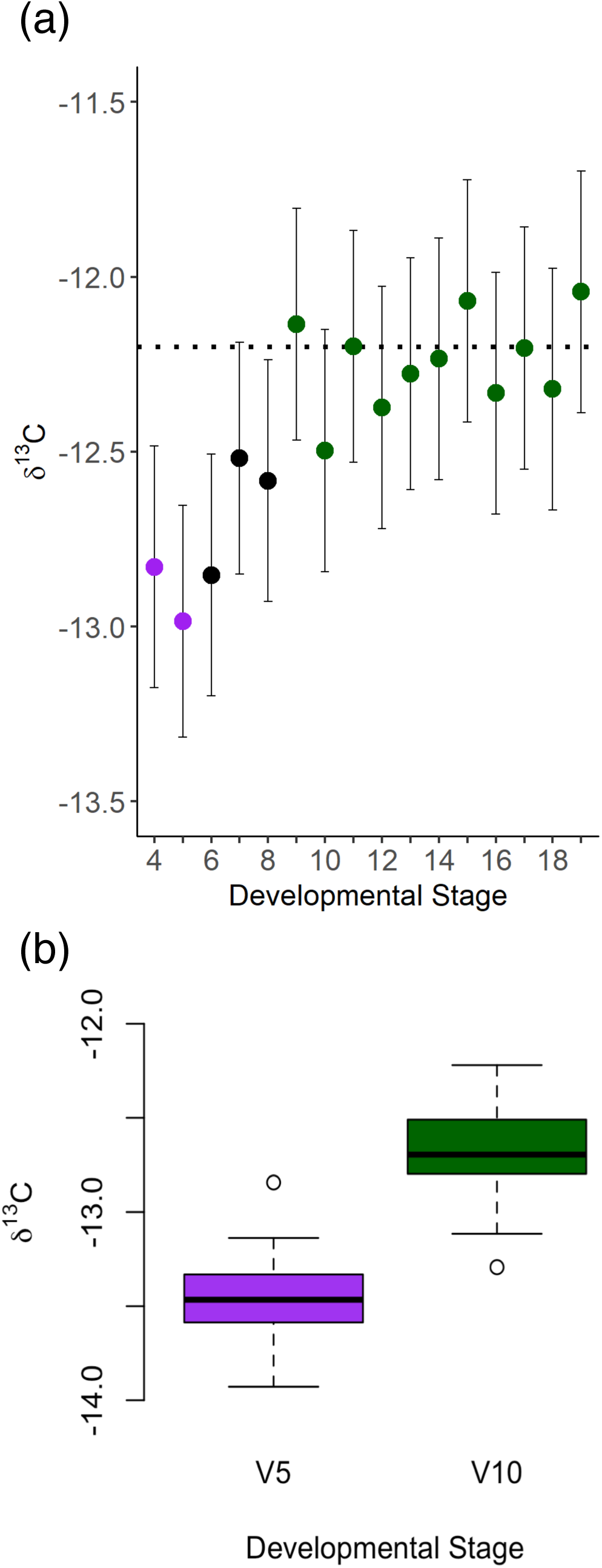
Developmental variation of leaf δ^13^C in *Z. mays*. (a) Data points represent Least Squares Mean of leaf δ^13^C values calculated for each developmental stage over two independent field seasons. V-stage is shown on the x-axis. Error bars show standard error and horizontal dashed line represents the average leaf δ^13^C value (−12.2 ‰) of mature leaf stages (V9-V19). Colors show the trend from juvenile (purple), thru transition (black), to adult (green). (b) The inset boxplots shows juvenile and adult leaf δ^13^C values for 31 *Z. mays* lines. The box 1^st^ and 3^rd^ quartile with horizontal lines indicating the mean and the whiskers showing the mild outliers. Open circles represent the extreme outliers.

To assess whether differences in leaf δ^13^C between juvenile and adult leaves was significant and common across lines of *Z. mays*, the 31 inbred lines previously grown to assess diversity were sampled at leaf stages V5 and V10. The juvenile δ^13^C leaf samples were taken at V5 to reduce the possibility of sampling leaves that are composed primarily of carbon from the kernel, which can contribute significantly to the first few leaves. Sampling later than V5 would have introduced the possibility of inadvertently measuring within the stages of transition (V6-V8). A highly significant difference in δ^13^C was observed between juvenile and adult leaves (*p* < 0.0001). The shift to less negative leaf δ^13^C values (average V5 = −13.56 ‰ and V10 = −12.67 ‰) were consistent with what was observed with B73 (Figure 3B). Although all 31 lines shifted in the same direction, the amount varied from 0.50 % to 1.30 % depending on the line, with an average of 0.79 ‰ across all lines (Figure S1). Significant differences in leaf percent carbon (*p* < 0.0001), percent nitrogen (*p* = 0.0090), and δ^15^N (*p* = 0.0009) were also observed between juvenile and adult leaves (Figure S2).

To further investigate the difference in δ^13^C observed between juvenile to adult leaves, samples were taken from wild-type and *glossy15* (*gl15*) mutants in two different *Z. mays* backgrounds (B73 and W64A). It is well known that *gl15* mutants have a reduced expression of juvenile leaf traits and show adult leaf characteristics at V3 (Moose and Sisco, 1994). Premature adult leaf traits include adult epidermal wax, adult cell wall characteristics, presences of epidermal hairs, and the presence of bulliform cells. Although only two replicates were grown for each genotype and were sampled as pools, no differences were observed between wild-type and *gl15* mutants in either background. At V5 the difference between wild-type and *gl15* mutant plants was −0.07 % and 0.02 % in B73 (average −13.03 ‰) and W64A (average −13.28 ‰) respectively. At V10 the difference between wild-type and *gl15* mutant plants was −0.06 % and −0.16 % in B73 (average - 12.87 ‰) and W64A (average −12.83 ‰) respectively. Because no obvious differences were observed between wild-type and *gl15* mutants, these experiments were not repeated. These results suggest that epidermal leaf characteristics under the control of *Gl15* are not responsible for the differences in leaf δ^13^C observed in juvenile and adult leaves.

### Relationship between Leaf δ^13^C and Water Stress in *Zea mays*

A subset of lines with consistently extreme leaf δ^13^C values were grown for three field seasons. This subset was made up of four Recombinant Inbred Lines (RILs) with high (less negative) leaf δ^13^C values and four RILs with low (more negative) leaf δ^13^C values. The high and low leaf δ^13^C RIL groups had distinct isotopic signatures when grown in 2015 and 2016 (Figure 4; *p* < 0.0001). Both the 2015 and 2016 growing season had ample rainfall from the time of planting until the time of sampling (23.5 cm and 25.8 cm, respectively). The consistent difference between the high and low RIL groups is not present in 2017 (Figure 4), when there was no significant difference between the high and low RIL groups (*p* = 0.0744). Interestingly, 2017 was a dry year with only 14.0 cm of precipitation between planting and sampling, and plants showed drought phenotypes for much of the growing season.

**Figure 4:**
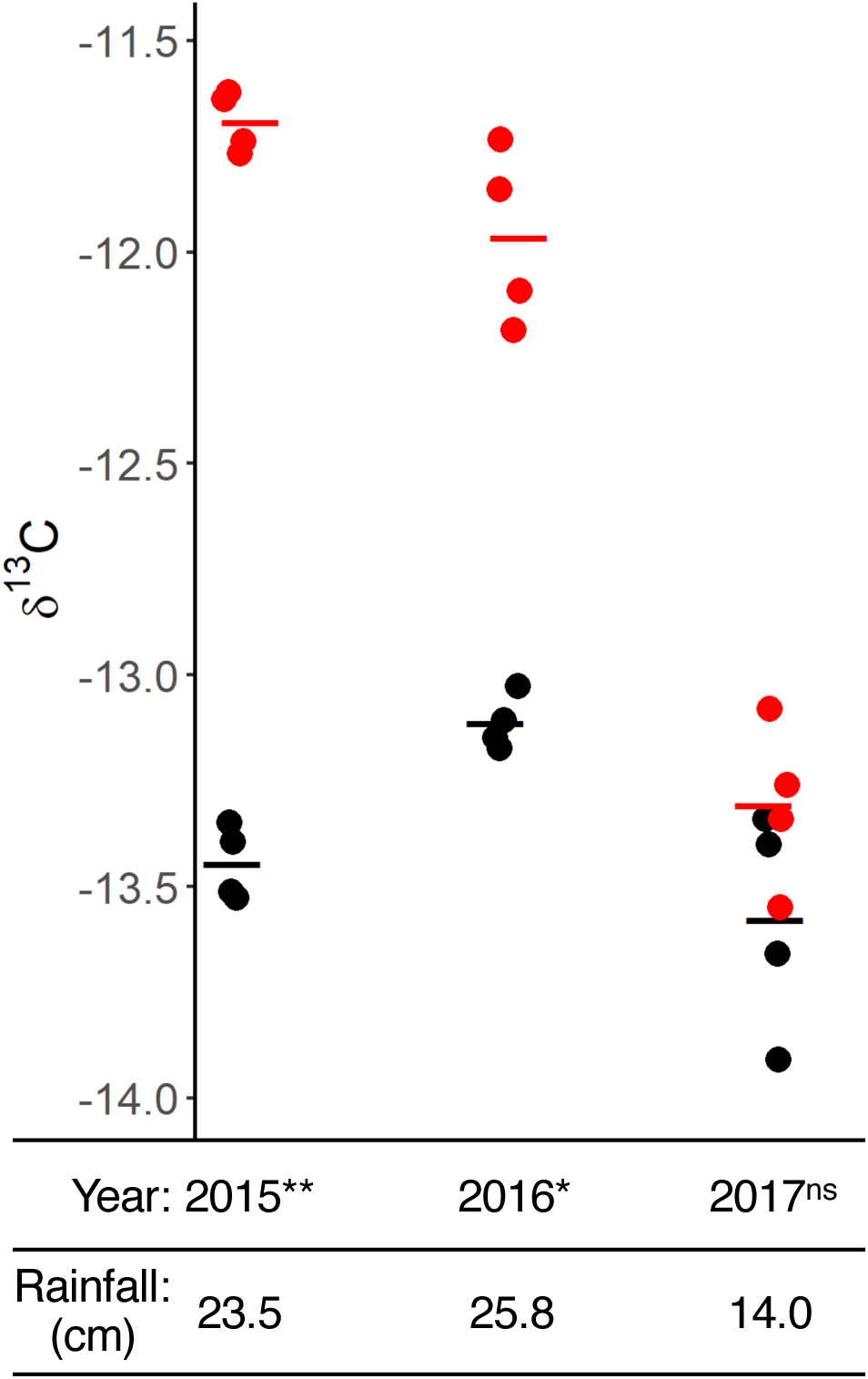
Response of leaf δ^13^C to water limiting conditions. Points represent leaf δ^13^C values of eight RILs grown in three consecutive field seasons. Horizontal lines indicate averages of the “high” (red) and “low” (black) RIL groups in each year. Rainfall totals (in centimeters) from the time of planting to sampling are shown below each year. ** *p* = 3.685 × 10^−8^, * *p* = 2.185 × 10^−5^, ns (not significant).

Because the results of the field experiments suggested a link between leaf δ^13^C and drought stress in *Z. mays*, a greenhouse experiment was performed to more precisely tease apart this relationship. Of the eight RILs grown in the field that had extreme isotope values, four were selected that had similar plant height and flowering time. The subset of RILs selected for the greenhouse experiment was comprised of two RILs with high leaf δ^13^C values (Z007E0067 and Z021E0032) and two RILs with low leaf δ^13^C values (Z007E0150 and Z021E0097), when measured in the field. To ascertain the effect of water availability on leaf δ^13^C, the four RILs were grown under three different water treatments: 100% field capacity (FC), 80% FC, and 40% FC.

A significant treatment effect was observed between 40% FC and both 100% and 80% FC with respect to the total amount of water transpired (*p* <0.00001; Figure 5). However, no difference in total transpiration was observed between the 100% and 80% FC treatments (*p* = 0.3547). A genotypic effect was observed with Z007E0067 transpiring significantly more than Z007E0150 at both 100% and 80% FC (*p* = 0.02527 and *p* = 0.0306 respectively; Figure 5, Figure S3). No significant genotypic difference was observed for total transpiration when plants were grown at 40% FC (*p* =0.2260). However, all four RILs grown at 40% FC had significantly lower leaf δ^13^C values (*p* <0.0001) than both the 100% and 80% FC treatments (Figure 5). A clear decrease in leaf δ^13^C is observed as total transpiration is reduced (Figure 5). A genotypic effect was also seen in regards to leaf δ^13^C. Z007E0067 had significantly higher leaf δ^13^C values than all other genotypes at 100% and 80% FC (*p* = 0.00044 and *p* = 0.00118 respectively; Table 1). For the 40% FC treatment, Z007E0150 and Z007E0067 had significantly higher δ^13^C values when compared to Z021E0097 (*p* = 0.00114 and *p* = 0.03994 respectively; Table 1).

**Figure 5:**
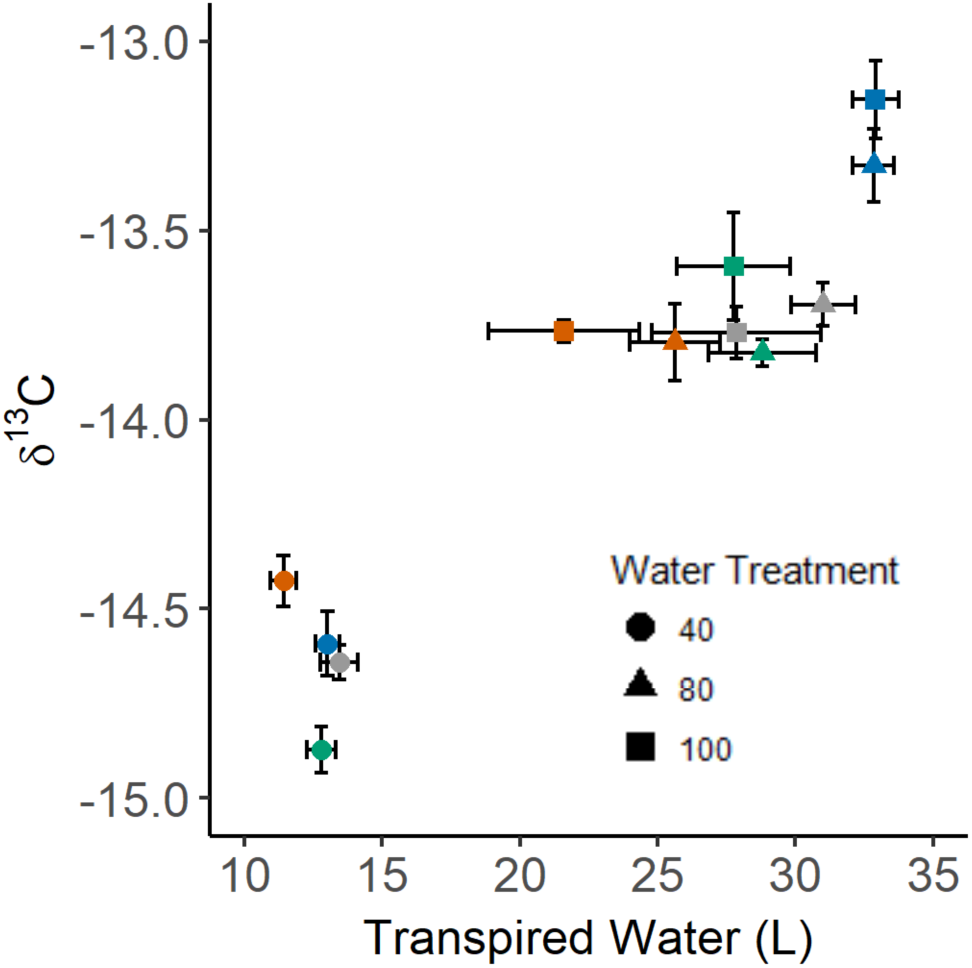
Leaf δ^13^C compared to transpiration of four *Z. mays* RILs grown under three water treatments. Gray (Z021E0032), orange (Z007E0150), green (Z021E0097), and blue (Z007E0067) points represent means of the RILs grown in each treatment. Bars show standard error of the mean. Shape of the points distinguish 100%, 80%, and 40% FC treatment. The treatment effect between 100% FC and 80% FC was not significant, but the treatment effect between 40% FC and both 80% FC and 100% FC was significant at *p* <0.00001.

**Table 1.** Gas Exchange Measurements of 100% FC Greenhouse Grown RILs. Presented as average ± standard error. Significance threshold of *p* < 0.05.

| Genotype | Net<br>Photosynthesis<br>(*A*) | Stomatal<br>conductance<br>(*g<sub>s</sub>*) | Transpiration<br>(*E*) | Intercellular<br>CO<sub>2</sub>/Atmospheric<br>CO<sub>2</sub> (*C<sub>i</sub>/C<sub>a</sub>*) | TE<sub>i</sub><br>(*A/E*) |
| --- | --- | --- | --- | --- | --- |
|  | μmol m<sup>-2</sup> s<sup>-1</sup> | mol m<sup>-2</sup> s<sup>-1</sup> | mol m<sup>-2</sup> s<sup>-1</sup> | μmol mol<sup>-1</sup> | μmol m<sup>-2</sup> s<sup>-1</sup><br>mol m<sup>-2</sup> s<sup>-1</sup> |
| Z007E0067 | 32.2<br>± 0.7<sup>a</sup> | 0.268<br>± 0.012<sup>a</sup> | 0.0039<br>± 0.0002<sup>a</sup> | 0.3852<br>± 0.0155<sup>a</sup> | 8406.86<br>± 252.15<sup>b</sup> |
| Z007E0150 | 27.2<br>± 0.5<sup>b</sup> | 0.185<br>± 0.004<sup>b</sup> | 0.0027<br>± 0.0001<sup>b</sup> | 0.2825<br>± 0.0130<sup>b</sup> | 10024.77<br>± 177.28<sup>a</sup> |
| Z021E0032 | 34.3<br>± 1.0<sup>a</sup> | 0.260<br>± 0.011<sup>a</sup> | 0.0038<br>± 0.0002<sup>a</sup> | 0.3215<br>± 0.0115<sup>b</sup> | 9151.14<br>± 200.11<sup>ab</sup> |
| Z021E0097 | 32.3<br>± 1.7<sup>a</sup> | 0.250<br>± 0.017<sup>a</sup> | 0.0036<br>± 0.0002<sup>a</sup> | 0.3419<br>± 0.0219<sup>ab</sup> | 8930.24<br>± 292.95<sup>b</sup> |

To investigate potential leaf level gas exchange differences between RILs, we collected steady state gas exchange measurements on all four RILs grown in the 100% FC treatment along with samples for leaf δ^13^C at 54 days after planting (DAP). Line Z007E0150 had significantly lower values of net photosynthesis (*A*), stomatal conductance (*g*_*s*_), and transpiration (*E*) compared to the other RILs (*p* <0.05; Table 2).In addition, Z007E0150 had the lowest *C*_*i*_/*C*_*a*_ value (98.7 ± 4.6), which was significantly lower than Z007E0067 (130.8 ± 5.1; *p* = 0.0021). Z021E0032 also had a *C*_*i*_/*C*_*a*_ significantly lower than Z007E0067 (*p* = 0.0266; Table 2). A comparison of leaf δ^13^C collected at 54 DAP (corresponding to the gas exchange measurements) and at 64 DAP (corresponding to the termination of the experiment) showed no significant differences in any of the four RILs (Figure S4). Consistent with field measurements, Z007E0067 had a significantly higher leaf δ^13^C; however, in the greenhouse Z021E0032 was statistically similar to the low leaf δ^13^C RILs (Table 1).

**Table 2.**
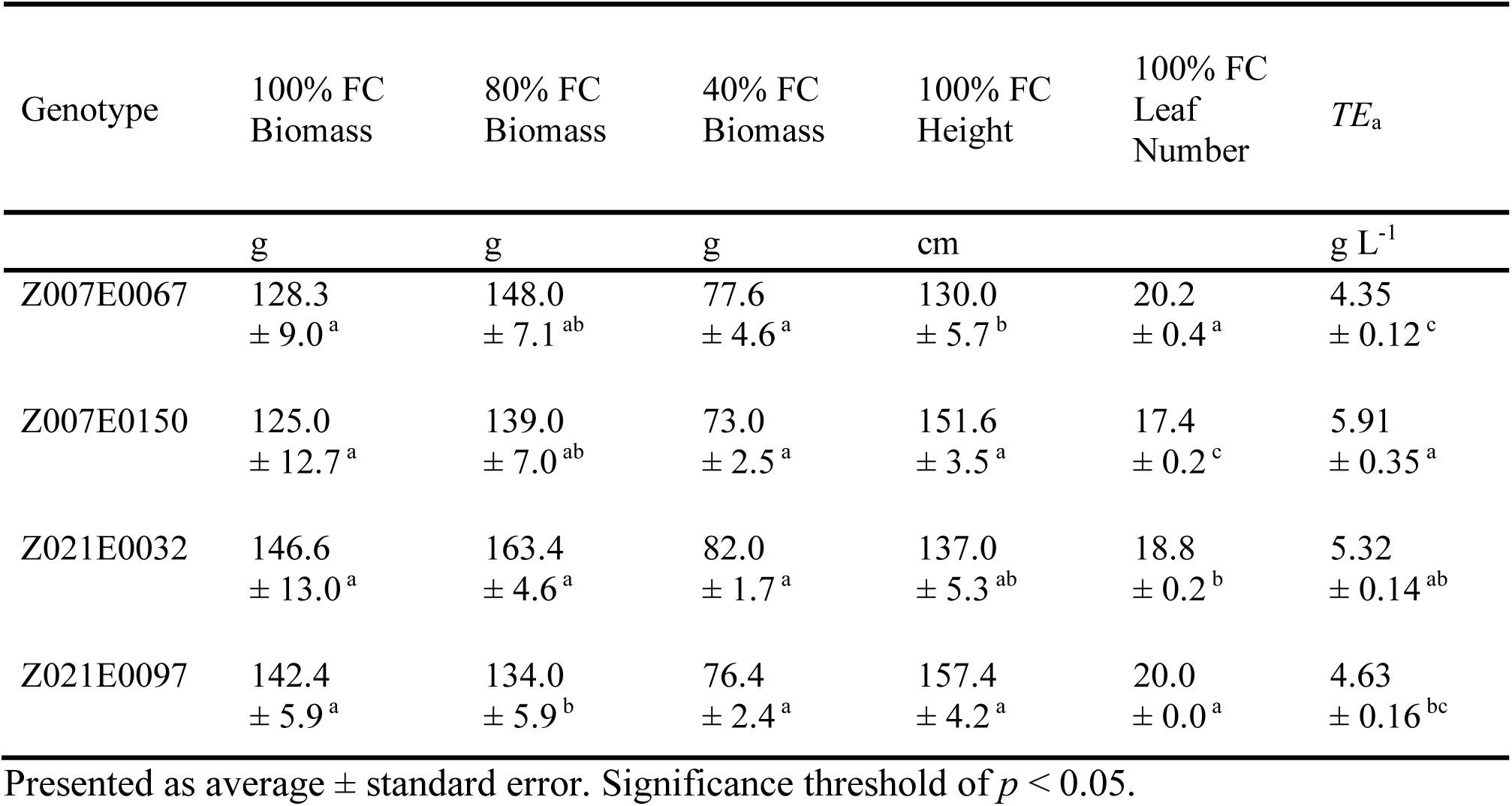
Agronomic Measurements of Greenhouse Grown RILs. Presented as average ± standard error. Significance threshold of *p* < 0.05.

| Genotype | 100% FC<br>Biomass | 80% FC<br>Biomass | 40% FC<br>Biomass | 100% FC<br>Height | 100% FC<br>Leaf<br>Number | *TE*<sub>a</sub> |
| --- | --- | --- | --- | --- | --- | --- |
|  | g | g | g | cm |  | g L<sup>-1</sup> |
| Z007E0067 | 128.3<br>± 9.0<sup>a</sup> | 148.0<br>± 7.1<sup>ab</sup> | 77.6<br>± 4.6<sup>a</sup> | 130.0<br>± 5.7<sup>b</sup> | 20.2<br>± 0.4<sup>a</sup> | 4.35<br>± 0.12<sup>c</sup> |
| Z007E0150 | 125.0<br>± 12.7<sup>a</sup> | 139.0<br>± 7.0<sup>ab</sup> | 73.0<br>± 2.5<sup>a</sup> | 151.6<br>± 3.5<sup>a</sup> | 17.4<br>± 0.2<sup>c</sup> | 5.91<br>± 0.35<sup>a</sup> |
| Z021E0032 | 146.6<br>± 13.0<sup>a</sup> | 163.4<br>± 4.6<sup>a</sup> | 82.0<br>± 1.7<sup>a</sup> | 137.0<br>± 5.3<sup>ab</sup> | 18.8<br>± 0.2<sup>b</sup> | 5.32<br>± 0.14<sup>ab</sup> |
| Z021E0097 | 142.4<br>± 5.9<sup>a</sup> | 134.0<br>± 5.9<sup>b</sup> | 76.4<br>± 2.4<sup>a</sup> | 157.4<br>± 4.2<sup>a</sup> | 20.0<br>± 0.0<sup>a</sup> | 4.63<br>± 0.16<sup>bc</sup> |

From this experiment both *TE*_i_ and *TE*_a_ were calculated at 100% FC. Intrinsic transpiration efficiency, calculated using gas exchange measurements, showed that Z007E0150 had a significantly higher *TE*_i_ compared to Z007E0067 and Z021E0097 (*p* = 0.0006, *p* = 0.0237; Table 3). Agronomic transpiration efficiency, calculated using biomass and total water transpired, also showed that RIL Z007E0150 had a significantly higher *TE*_a_ than Z007E0067 and Z021E0097 (*p* = 0.0006 and *p* = 0.0059 respectively; Table 3). Z021E0032 had significantly higher *TE* than Z007E0067, but only when calculating *TE*_a_ (*p* = 0.0262). To further investigate what was driving *TE*_a_ differences between RIL lines, we looked closer at plant biomass. No significant difference in biomass was found between RILs at 100% FC (*p* = 0.43), indicating that the difference in *TE*_a_ are mainly attributable to total transpiration. However, significant differences were observed in plant height and leaf number (Table 3). Most notably, Z007E0150 is one of the taller RILs but has significantly fewer leaves, whereas Z007E0067 is the shortest RIL and has the most leaves (Table 3).

**Table 3.** Stable Carbon Isotope Values of Greenhouse Grown RILs. Presented as average ± standard error. Significance threshold of *p* < 0.05.

| Genotype | 100% FC<br>54 DAP ($\delta^{13}\text{C}$) | 100% FC<br>64 DAP ($\delta^{13}\text{C}$) | 80% FC<br>64 DAP ($\delta^{13}\text{C}$) | 40% FC<br>64 DAP ($\delta^{13}\text{C}$) |
| --- | --- | --- | --- | --- |
|  | ‰ | ‰ | ‰ | ‰ |
| Z007E0067 | $-13.18 \pm 0.03^a$ | $-13.15 \pm 0.10^a$ | $-13.33 \pm 0.10^a$ | $-14.59 \pm 0.09^a$ |
| Z007E0150 | $-13.64 \pm 0.05^b$ | $-13.77 \pm 0.03^b$ | $-13.80 \pm 0.10^b$ | $-14.43 \pm 0.07^a$ |
| Z021E0032 | $-13.60 \pm 0.02^b$ | $-13.77 \pm 0.07^b$ | $-13.70 \pm 0.06^b$ | $-14.64 \pm 0.05^{ab}$ |
| Z021E0097 | $-13.70 \pm 0.04^b$ | $-13.60 \pm 0.14^b$ | $-13.82 \pm 0.04^b$ | $-14.87 \pm 0.06^b$ |

Presented as average ± standard error. Significance threshold of $p < 0.05$ .

To determine the correlation between traits measured during the greenhouse experiment, a correlation matrix was generated using traits collected in all three water treatments (Figure 6). As expected, strong negative correlations were found between total transpiration and *TE*_a_ as well as between biomass and *TE*_a_ because both total transpiration and biomass are used to calculate *TE*_a_. However, a negative correlation was also observed between *TE*_a_ and δ^13^C values. Large positive correlations were found between leaf δ^13^C values and total transpiration, leaf δ^13^C values and biomass, and total transpiration and biomass.

**Figure 6:**
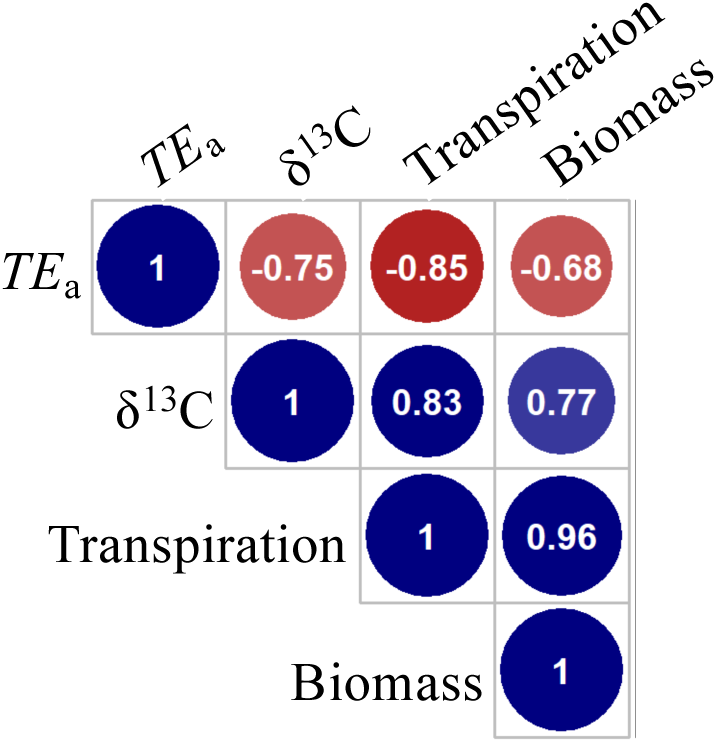
Pearson coefficient for phenotypic traits recorded for all water treatments. Blue shades represent positive correlations while red shades represent negative correlations. The size of the circles represent the strength of correlation with larger circles representing stronger correlations.

## Discussion

Despite the complexity of isotopic fractionation in C_4_ species, the use of leaf δ^13^C has been proposed as a method for measuring *TE* in *Z. mays*. Here we specifically test field-grown *Z. mays* and sources of variation in sampling leaf δ^13^C. These results facilitate the standardization and selection of the appropriate sampling time to maximize the effectiveness of leaf δ^13^C measurements. Furthermore, we specifically test the relationship between *TE* and leaf δ^13^C in the greenhouse. These results open up specific targets for the optimizing of water-use efficiency in a major C_4_ crop.

The 31 inbred lines of *Z. mays* sampled in this study had a trait distribution for leaf δ^13^C that spans a large portion of the range seen across all C_4_ species (−11 to −15 ‰; O’Leary, 1988). Because the observed variation has a moderate genetic heritability, this trait has the potential to be used in a breeding program. The heritability for leaf δ^13^C is similar to what has previously been reported in *Z. mays* (0.51; Foley, 2012). Because our goal was to assess natural genetic variation in leaf δ^13^C, the lines sampled were selected to maximize diversity. Therefore, only two ExPVP lines were included in this study. Thus, it is not possible with this data set to assess the amount of variability for leaf δ^13^C found in temperate elite germplasm compared to the diverse NAM founder lines. However, both of the ExPVP lines were found to have values at the high (less negative) end of the trait distribution, possibly indicating that there are alleles from diverse material that could lower leaf δ^13^C values.

Similar variation in leaf δ^13^C was reported for the NAM Founder lines when grown in the greenhouse (Kolbe *et al.*, 2017). Although many of the lines were stable in their relative position in the distribution, the absolute value of the trait distribution seen in the greenhouse was shifted to more negative leaf δ^13^C values. While there are many differences between greenhouse and field-grown plants, this shift is likely due to the differences in the atmospheric concentration of ^13^C between greenhouse (−11 ‰; Kolbe *et al.*, 2017) and field (typically −8 ‰) environments.

The shift to more negative leaf δ^13^C values is also visible in our greenhouse experiment when compared to field-grown plants of the same RILs. While most of the lines were relatively consistent between greenhouse and field environments, some lines had significant genotype by environment interactions as seen by the drastic change of their relative rank. In particular, CML247 and W22 had high leaf δ^13^C values in the greenhouse but consistently low leaf δ^13^C in the field. The opposite was true for CML228. A genotype by environment interaction was also observed with RIL Z021E0032. This would suggest that greenhouse screens of leaf δ^13^C are useful, but would need to be supported with field experiments to confirm the phenotype.

The investigation of variation in sampling with respect to circadian and developmental time revealed periods of stable leaf δ^13^C. Here we performed a formal testing of the hypothesis that leaf δ^13^C is an integrated measurement of fixed carbon rather than an instantaneous measure of the active photosynthetic pool. The relative stability of leaf δ^13^C over the circadian time-course experiment indicates the time at which the sample is taken during the day has no effect on the isotopic signature. Unlike the stability seen over a 24-hour period, there was a significant trend in leaf δ^13^C over developmental time. Over multiple years and across a diverse panel of lines, there was a significant difference between leaf δ^13^C in juvenile and adult leaves. The timing of transition from low to high leaf δ^13^C seen in this study is similar to what was measured previously in a coarse developmental time series (Zhang *et al.*, 2015). Although the difference in leaf δ^13^C through development was not as large as between genotypes, an average difference of % was observed between V5 and V10 leaves (Figure 2B).

At this point it is not clear what process is driving the difference in leaf δ^13^C between juvenile and adult leaves. Interestingly, the timing of the change is similar to vegetative phase change timing in *Z. mays*. An obvious indicator of this transition is the change from juvenile to adult epicuticular wax, which occurs around V7 in B73 (Foerster *et al*, 2015). Due to the possible connection, differences in vegetative phase change timing between lines should be considered when sampling. However, >95% of the 4,018 NAM RILs tested transitioned before V10 (Foerster *et al*, 2015), which supports our conclusion that V10 is near optimal for sampling leaf δ^13^C. Although later leaf stages are stable with respect to leaf δ^13^C, other differences between lines are compounded as the growing season progresses. Our analysis of *gl15* mutants suggests that the difference in leaf δ^13^C is not due to changes in epicuticular wax. In addition to changes in epicuticular wax, several other leaf characteristics change during the juvenile to adult phase change including thickening of the cuticle, changes in epidermal cell walls, presences of epidermal hairs, and the presence of bulliform cells (Poethig, 1990; Moose and Sisco, 1994). The effect of cuticle thickness on leaf δ^13^C is still not known because it is not controlled by *Gl15* (Evans *et al.*, 1994). In addition to leaf morphology, it is also possible that the difference in leaf δ^13^C is due to physiological differences between juvenile and adult leaves. This could include differences in stomatal opening that could cause a low *C*_*i*_*/C*_*a*_ in juvenile leaves. Alternatively, the difference in leaf δ^13^C may not be due to characteristics of the juvenile leaf per se, but rather it could be due to the carbon signature from the previous generation, which contributes significantly to juvenile leaves through seed reserves.

Previous reports on the effect of drought on leaf δ^13^C in *Z. mays* have been inconsistent (Monneveux et al., 2007; Zhang *et al.*, 2015). Theoretically, different trends are possible if Φ significantly changes under drought stress. When Φ is > 0.37, a negative relationship between δ^13^C and *C*_i_/*C*_a_ is predicted; however, if Φ is < 0.37, a positive relationship between δ^13^C and *C*_i_/*C*_a_ is predicted (Cernusak *et al.* 2013). Although estimates of Φ from gas exchange measurements and dry leaf δ^13^C have documented substantial changes in Φ in response to environmental differences, the more accurate measure of Φ calculated from instantaneous measurements of carbon isotope discrimination is relatively constant under different irradiances, temperatures, and CO_2_ concentrations (Kromdijk *et al.*, 2014). However, because there is a lack of direct measurements of Φ under water limiting conditions, the extent to which Φ influences leaf δ^13^C under these conditions is not known. Our data show a clear decrease in leaf δ^13^C under water limiting conditions for field-grown plants as well as plants grown under controlled water stress experiments in the greenhouse. This result is consistent with a Φ < 0.37 where the reduction in *C*_*i*_/*C*_*a*_ in response to water limitations is the major driver of leaf δ^13^C under drought. A similar result was found in the C_4_ grasses *Setaria viridis* and *Setaria italica* (Ellsworth *et al.*, 2017). However, our data from the greenhouse experiment indicates that while *C*_*i*_/*C*_*a*_ may be a major factor, it is not the only thing effecting leaf δ^13^C, as seen by directly comparing *C*_*i*_/*C*_*a*_ to leaf δ^13^C (Table 1 and 2). Other factors such as mesophyll conductance could also be contributing to leaf δ^13^C, but it is predicted that the effect of mesophyll conductance on leaf δ^13^C in C_4_ plants is small (von Caemmerer *et al*., 2014).

The response of leaf δ^13^C to water limitation in the field, and the greenhouse, has several implications for leaf δ^13^C use as a proxy trait for water-use efficiency. Under severe drought it is possible that *C*_*i*_/*C*_*a*_ approaches a biological limit that drives leaf δ^13^C values to be more negative and nearly uniform across all lines. If there is a lower limit to *C*_*i*_/*C*_*a*_, it is interesting to speculate that the small differences observed in leaf δ^13^C between lines grown under drought conditions may reflect the amount of variation in other processes (such as Φ or post photosynthetic fractionation) that contribute to leaf δ^13^C. Regardless, these results seem to suggest that measurements of leaf δ^13^C may only be informative in assessing *TE* under well-watered conditions. Under 100% FC in the greenhouse, Z007E0067 had the highest transpiration and also had a significantly higher leaf δ^13^C compared to the other RILs. Thus, selection for leaf δ^13^C under well-watered conditions may provide a breeding target for improving *TE* and the sustainability of *Z. mays* production under current and future growing environments.

## Experimental Procedures

### Plant Growth

All field trials were grown at the Crop Sciences Research and Education Center located in Urbana, Illinois. Fifteen kernels were planted in each 3.7 meter row with 0.8 meter spacing between rows and 0.9 meter alleys. The fields were planted on May 14^th^ 2015, May 7^th^ 2016, and May 16^th^ 2017. No irrigation was provided for any of the experiments. All 31 *Z. mays* inbred lines used in these experiments are publicly available through the USDA Germplasm Resources Information Network (GRIN). These inbreds were planted as a single block arranged according to flowering time in 2015 and 2016. The *glossy15-H* allele used in this study was previously described by Moose and Sisco (1996), and were planted in two single row plots in 2016. Plants were sampled at the developmental stage indicated for each experiment.

The RIL greenhouse experiment was conducted at the University of Illinois Plant Care Facility located in Urbana, Illinois. NAM RILs used in this experiment are available from the Maize Genetics Stock Center. Kernels were sown in 50 cell trays filled with LC1 soilless media (SunGro Sunshine Mix #1). Plants were grown in the greenhouse under supplemental metal halide lighting set to 14 hour days. Prior to transplanting into Classic^®^ 2000 pots, a coffee filter was placed at the bottom of each pot to prevent any media from being lost and each pot was filled with the same volume of media. The media used was 3:1:1:1 of LC1, sterilized field soil, peat, and perlite respectively. The field capacity of the pots was determined by saturating the soil for 3 days and weighing each pot once the excess water had run through the media. The pots were then dried in a drier with continuous airflow at 60°C. Pots were periodically weighed until their weight remained constant thus signifying complete dry down, which occurred after 15 days. The difference for each pot between saturated soil and dry down was determined to be the weight of available water. This difference was averaged to determine the amount of water at field capacity (FC). The weight of 80% and 40% FC was calculated for each water treatment. Seedlings were transplanted into pots 17 days after planting (DAP) and the pots were arranged in a randomized complete block design. Soil amendments for fertility were added to the pots during transplanting which included: Osmocote^®^ (11-4-17), dolomitic lime, phosphorous, magnesium sulfate, and gypsum. Plants were well watered until the start of the experiment. The fungicide Pagent^®^ was applied directly after transplanting. For the duration of the experiment the pots were fertilized with 500 mL of 300 ppm (15-5-15) CalMag every 7 days. Before the water treatments began, each pot was weighed with the 21 day seedling and the treatment weight was calculated. The water treatment began 33 DAP. Each morning at the beginning of the light period, the pots were weighed and the difference between the previous day’s weight was calculated and the weight lost through evapotranspiration was replaced with the equivalent weight of water. In each water treatment replication there was a pot without a plant which served as an evaporation control. This allowed daily transpiration of each plant to be measured over the course of the treatment period (Figure S3). The experiment was terminated 64 DAP and leaf samples were taken from the upper most fully expanded leaf for isotopic analysis. Aboveground biomass was taken by cutting the plant at the soil surface and drying it at 60°C for 14 days. Total water use was calculated by summing the difference in weight each day and subtracting the difference in weight of the evaporation control pot per each treatment. *TE*_*a*_ was calculated by the aboveground biomass (g) over the total volume of water used during the treatment (L).

Gas exchange measurements were taken on the uppermost fully expanded leaf using a LICOR-6800 over three days starting 50 DAP. The uppermost fully expanded leaf was leaf 9 (V9) in all measured plants. Only plants from the 100% FC treatment group were measured. Five plants from each line were measured; with the exception of Z021E0097, only four plants were measured due to the loss of one replicate. All leaf chamber conditions were set constant during measurements. The temperature of the exchanger was set at 25°C and H_2_O concentrations were controlled automatically to maintain a constant VPD of 1.5 kPa. The CO_2_ concentration of the chamber was set constant at 400 µmol mol^−1^ for the duration of measurements and light was set at 1,500 µmol m^−2^ s^−1^. Each plant was acclimated to the chamber conditions for 15-30 minutes until steady state was achieved. Once the leaf reached steady-state, data was logged every 60 seconds for a duration of 5 minutes. Average net photosynthesis (*A*), stomatal conductance (*g*_*s*_), and transpiration (*E*) were calculated for each line and an ANOVA and Tukey’s *post hoc* tests were ran to determine significant differences. Intrinsic transpiration efficiency was calculated using the equation (*A*/*E*) and an ANOVA with Tukey’s *post hoc* test was run to determine significant difference between lines. Leaf samples for isotopic analysis were taken from the portion of the leaf that was measured via the Licor 54 DAP.

### Leaf Sample Collection

All samples for isotope analysis were collected from the middle of the leaf blade (avoiding the midrib) of the uppermost fully expanded leaf using a 0.5 cm diameter single hole punch (Office Depot #825232). Only healthy leaf tissue was sampled to avoid confounding factors. Leaf samples were collected from individual plants or as pools of multiple plants from a single row depending on the experiment. Individual samples consisted of 12-24 punches total, half taken from each side of the midrib from the same leaf of a single plant. Pooled samples consisted of 6 punches from four individual plants from a single row. Leaf punches were placed in 2 mL Simport tubes with screw caps (VWR # 89499-598). Three 1/8″ Grade 1000 Type 430 Stainless Steel ball bearings (Abbott Ball Company) were added to each tube for grinding. After collection, tubes containing tissue samples were uncapped and placed in a dryer at 65°C for at least 7 days. Tubes were then capped and placed in a Nalgene desiccation cabinet (VWR # 24987-056) containing Drierite until the samples were prepared for isotope analysis. For the circadian time-course experiment, samples were taken from three randomly selected individuals at each time point from a large block of B73 plants. Plants were only sampled once during the experiment, and thus the data represents 72 individual plants. The δ^13^C of three plants were measured individually and also as a combined pool.

### Stable Carbon Isotope Analysis

Dry leaf tissue was ground in the Simport tubes with ball bearings by shaking in a Geno Grinder at 1000 rpm for 10 minutes. Ground leaf tissue was weighed out into 6 mm × 4 mm tin capsules (OEA Laboratories # C11350.500P) using a Sartorius CPA2P scale. The range of acceptable sample weight varied between isotope facilities. Samples were placed in a Costar 96-well plate and stored in the desiccation cabinet until it was analyzed. Samples were run through a Costech instruments elemental combustion system and then either a Delta V Advantage (University of Illinois) or a Delta PlusXP (Washington State University) isotope ratio mass spectrometer to determine δ^13^C values. The precision of the instruments measuring δ^13^C is 0.2 ‰. Samples for the Diversity (2015), Circadian time-course (2016), B73 Development (2015), and greenhouse experiments were analyzed at The University of Illinois Urbana-Champaign. Samples for the Diversity (2016), B73 Development (2016), Juvenile/Adult leaves (2016), *gl15* experiment, and RILs (2015, 2016, 2017) were analyzed at Washington State University. The use of Vienna Peedee Belemnite calibrated standards permits the direct comparison of leaf δ^13^C from different facilities.

### Statistical Analyses

All statistical tests and data visualizations were performed using statistical packages in R (R Development Core Team, 2010). Graphs were made using the *boxplot* function or using the package *ggplot2*. A linear model was used for the B73 Development experiment to combine the data from 2015 and 2016 using the package *lme4*. Least Squares means were then calculated using the package *lsmeans*. One way ANOVAs were used to test for significant differences between data points in the B73 Developmental experiment, the Circadian time-course, and greenhouse experiment. Tukeys HSD with an alpha of 0.05 was performed using the package *multcomp* after each significant ANOVA test. A Pearson correlation test was performed to calculate the linear dependence of the response variables to each other. The correlation matrix was constructed using the package *corrplot*. Heritability of leaf δ^13^C was calculated using NAM Founder data from 2015 and 2016 described in this study. Data from Kolbe *et. al.*, (2017) was also used in the heritability calculation, which included three replicates of each of the NAM founder lines and six replicates of the reference line B73 grown in the greenhouse. Therefore, the calculation included three environments and 27 genotypes. First the variance components were estimated using *lme4*. Then broad sense heritability on a line mean basis was calculated using the equation

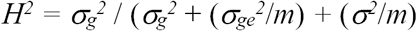

where 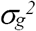 is the genotypic variance, 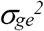 is the genotype:environement variance, *σ*^*2*^ is the error variance, and *m* is the number of environments, as described previously Piepho and Möhring, 2007).

## Acknowledgements

We thank Dr. Steve Moose for providing seed for the *glossy15* experiment. We would also like to thank Mengqiao Han for her technical help in isotope preparation and thoughtful discussion during manuscript preparation. The authors have no conflict of interest to declare. This work was supported by United States Department of Agriculture through Hatch funds.

## References

Blum, A. (2009) Effective use of water (EUW) and not water-use efficiency (WUE) is the target of crop yield improvement under drought stress. Field Crops Res., 112, 119–123.

Brüggemann, N., Gessler, A., Kayler, Z., Keel, S.G., Badeck, F., Barthel, M., Boeckx, P., Buchmann, N., Brugnoli, E., Esperschütz, J., and Gavrichkova, O. (2011) Carbon allocation and carbon isotope fluxes in the plant soil-atmosphere continuum: a review. Biogeosciences, 8, 3457–3489.

Cernusak, L.A., Ubierna, N., Winter, K., Holtum, J.A., Marshall, J.D., and Farquhar, G.D. (2013) Environmental and physiological determinants of carbon isotope discrimination in terrestrial plants. New Phytol., 200, 950–965.

Condon, A.G., Richards, R.A., Rebetzke, G.J., and Farquhar, G.D. (2004) Breeding for high water-use efficiency. J. Exp. Bot., 55, 2447–2460.

Dhanapal, A.P., Ray, J.D., Singh, S.K., Hoyos-Villegas, V., Smith, J.R., Purcell, L.C., King, C.A., Cregan, P.B., Song, Q., and Fritschi, F.B. (2015) Genome-wide association study (GWAS) of carbon isotope ratio (δ13C) in diverse soybean [Glycine max (L.) Merr.] genotypes. Theor. Appl. Genet., 128, 73–91.

Ellsworth, P.Z., and Cousins, A.B. (2016) Carbon isotopes and water use efficiency in C_4_ plants. Curr. Opin. Plant Biol., 31, 155–161.

Ellsworth, P.Z., Ellsworth, P.V., and Cousins, A.B. (2017) Relationship of leaf oxygen and carbon isotopic composition with transpiration efficiency in the C_4_ grasses *Setaria viridis* and *Setaria italica*. J. Exp. Bot., 68, 3513–3528.

Evans, M.M., Passas, H.J., znd Poethig, R.S. (1994) Heterochronic effects of *glossy15* mutations on epidermal cell identity in maize. Genes Devel., 120, 1971–1981.

Farquhar, G.D. (1983) On the nature of carbon isotope discrimination in C_4_ species. Funct. Plant Biol., 10, 205–226.

Farquhar, G.D., and Richards, R.A. (1984) Isotopic composition of plant carbon correlates with water-use efficiency of wheat genotypes. Funct. Plant Biol., 11, 539–552.

Farquhar, G.D., Ehleringer, J.R., and Hubick, K.T. (1989) Carbon isotope discrimination and photosynthesis. Ann. Rev. Plant Biol., 40, 503–537.

Foerster, J.M., Beissinger, T., de Leon, N., and Kaeppler, S. (2015) Large effect QTL explain natural phenotypic variation for the developmental timing of vegetative phase change in maize (*Zea mays* L.). Theor. Appl. Genet., 128, 529–538.

Foley R.C. (2012) The genetic diversity of water use efficiency in the nested association mapping population of Zea mays. Masters thesis, Department of Horticulture and Landscape Architecture, Purdue University.

Flint-Garcia, S.A., Thuillet, A.C., Yu, J., Pressoir, G., Romero, S.M., Mitchell, S.E., Doebley, J., Kresovich, S., Goodman, M.M., and Buckler, E.S. (2005) Maize association population: a high-resolution platform for quantitative trait locus dissection. Plant J., 44, 1054–1064.

Hall, A.J., and Richards, R.A. (2013) Prognosis for genetic improvement of yield potential and water-limited yield of major grain crops. Field Crops Res., 143, 18–33.

Hansey, C.N., Johnson, J.M., Sekhon, R.S., Kaeppler, S.M., and Leon, N.D. (2011) Genetic diversity of a maize association population with restricted phenology. Crop Sci., 51, 704–715.

Henderson, S.A., Caemmerer, S.V., and Farquhar, G.D. (1992) Short-term measurements of carbon isotope discrimination in several C_4_ species. Funct. Plant Biol., 19, 263–285.

Kim, T.H., Böhmer, M., Hu, H., Nishimura, N., and Schroeder, J.I. (2010) Guard cell signal transduction network: advances in understanding abscisic acid, CO_2_, and Ca_2_^+^ signaling. Ann. Rev. Plant Biol., 61, 561–591.

Kolbe A.R., Studer A.J., Cousins A.B. (2017) Biochemical and transcriptomic analysis of maize diversity to elucidate drivers of leaf carbon isotope composition. Funct. Plant Biol., FP17265

Kromdijk, J., Ubierna, N., Cousins, A.B., and Griffiths, H. (2014) Bundle-sheath leakiness in C_4_ photosynthesis: a careful balancing act between CO_2_ concentration and assimilation. J. Exp. Bot, 65, 3443–3457.

Lobell, D.B., Roberts, M.J., Schlenker, W., Braun, N., Little, B.B., Rejesus, R.M., and Hammer, G.L. (2014) Greater sensitivity to drought accompanies maize yield increase in the US Midwest. Science, 344, 516–519.

Long, S.P., and Bernacchi, C.J. (2003) Gas exchange measurements, what can they tell us about the underlying limitations to photosynthesis? Procedures and sources of error. J. Exp.Bot., 54, 2393–2401.

McMullen, M.D., Kresovich, S., Villeda, H.S., Bradbury, P., Li, H., Sun, Q., Flint-Garcia S., Thornsberry J., Acharya C., Bottoms C., and Brown, P. (2009) Genetic properties of the maize nested association mapping population. Science, 325, 737–740.

Monneveux, P., Sheshshayee, M.S., Akhter, J., and Ribaut, J.M. (2007) Using carbon isotope discrimination to select maize (*Zea mays* L.) inbred lines and hybrids for drought tolerance. Plant Sci., 173, 390–396.

Moose, S.P., and Sisco, P.H. (1994) *Glossy15* controls the epidermal juvenile-to-adult phase transition in maize. Plant Cell, 6, 1343–1355.

Moose, S.P., and Sisco, P.H. (1996) *Glossy15*, an APETALA2-like gene from maize that regulates leaf epidermal cell identity. Genes Devel. 10, 3018–3027.

O’Leary, M.H. (1981) Carbon isotope fractionation in plants. Phytochemistry, 20, 553–567.

O’Leary, M.H. (1988) Carbon isotopes in photosynthesis. Bioscience, 38, 328–336.

Piepho, H. P., and Möhring, J. (2007) Computing heritability and selection response from unbalanced plant breeding trials. Genetics, 177, 1881–1888.

Poethig, R.S. (1990) Phase change and the regulation of shoot morphogenesis in plants. Science, 250, 923–930.

Pryor, S.C., Barthelmie, R.J., and Schoof, J.T. (2013) High-resolution projections of climate-related risks for the Midwestern USA. Climate Res., 56, 61–79.

R Development Core Team, (2010) R: A language and environment for statistical computing. Vienna, Austria: R Foundation for Statistical Computing. Available at: https://www.r-project.org/.

Rebetzke, G.J., Condon, A.G., Richards, R.A., and Farquhar, G.D. (2002) Selection for reduced carbon isotope discrimination increases aerial biomass and grain yield of rainfed bread wheat. Crop Sci., 42, 739–745.

Romay, M.C., Millard, M.J., Glaubitz, J.C., Peiffer, J.A., Swarts, K.L., Casstevens, T.M., Elshire R.J., Acharya C.B., Mitchell S.E., Flint-Garcia S.A., and McMullen, M.D. (2013) Comprehensive genotyping of the USA national maize inbred seed bank. Genome Biol., 14, R55.

Saranga, Y., Jiang, C.X., Wright, R.J., Yakir, D., and Paterson, A.H. (2004) Genetic dissection of cotton physiological responses to arid conditions and their inter-relationships with productivity. Plant Cell Environ., 27, 263–277.

Teulat, B., Merah, O., Sirault, X., Borries, C., Waugh, R., and This, D. (2002) QTLs for grain carbon isotope discrimination in field-grown barley. Theor. Appl. Genet., 106, 118–126.

Tian, F., Stevens, N.M., and Buckler, E.S. (2009) Tracking footprints of maize domestication and evidence for a massive selective sweep on chromosome 10. Proc. Nat. Acad. Sci., 106, 9979–9986.

von Caemmerer, S. (2000) Biochemical models of leaf photosynthesis. CSIRO Publishing, Collingwood, Australia

von Caemmerer, S., Ghannoum, O., Pengelly, J.J., and Cousins, A.B. (2014) Carbon isotope discrimination as a tool to explore C_4_ photosynthesis. J. Exp. Bot., 65, 3459–3470.

Wallace, J.G., Larsson, S.J., and Buckler, E.S. (2014) Entering the second century of maize quantitative genetics. Heredity, 112, 30.

Zhang, C., Zhang, J., Zhang, H., Zhao, J., Wu, Q., Zhao, Z., and Cai, T. (2015) Mechanisms for the relationships between water-use efficiency and carbon isotope composition and specific leaf area of maize (*Zea mays* L.) under water stress. Plant Growth Regulation, 77, 233–243.

